# A three-dimensional thalamocortical dataset for characterizing brain heterogeneity

**DOI:** 10.1101/2020.05.22.111617

**Authors:** Judy A. Prasad, Aishwarya H. Balwani, Erik C. Johnson, Joseph D. Miano, Vandana Sampathkumar, Vincent de Andrade, Kamel Fezzaa, Ming Du, Rafael Vescovi, Chris Jacobsen, Konrad P. Kording, Doga Gürsoy, William Gray-Roncal, Narayanan Kasthuri, Eva L. Dyer

## Abstract

Neural cytoarchitecture is heterogeneous, varying both across and within brain regions. The consistent identification of regions of interest is one of the most critical aspects in examining neurocircuitry, as these structures serve as the vital landmarks with which to map brain pathways. Access to continuous, three-dimensional volumes that span multiple brain areas not only provides richer context for identifying such landmarks, but also enables a deeper probing of the microstructures within. Here, we describe a three-dimensional X-ray microtomography imaging dataset of a well-known and validated thalamocortical sample, encompassing a range of cortical and subcortical structures. In doing so, we provide the field with access to a micron-scale anatomical imaging dataset ideal for studying heterogeneity of neural structure.

## Background and Summary

Whether focusing on a large swath of cortex or a single subcortical nucleus, consistent and reliable visualization of cytoarchitecture is critical for the creation of reference points which demarcate the brain’s landscape [1]. This is true not only for the identification of landmarks (or regions of interest), but also the study of local circuits therein. Indeed, it is the distinguishing features in brain cytoarchitecture which arise at small, local scales (i.e., through clusters of cells which are packed in discrete barrels or layers; see [2, 3]) that continue to emerge across larger spatial scales to reveal the presence of functionally distinct regions. Thus, detailed views into the brain’s architecture can be used to experimentally manipulate circuits, and to advance the field’s understanding and integration of each of these overarching systems.

With advances in the reconstruction and analysis of significantly larger brain volumes, neuroscientists are now able to visualize patterns of microarchitecture that arise at a scale previously inaccessible using traditional methods [4, 5, 6]. Examples such as CLARITY [7], expansion microscopy [8], serial two photon tomography [9, 10, 11], multi-beam scanning electron microscopy [12], and X-ray microtomography [13, 14, 15, 16, 17], now provide access to several regions of interest within a volume of tissue simultaneously, providing rich context to study both local circuitry and long-range projections. With many of these new techniques, it is possible to image and analyze large intact anatomical samples that preserve the connectivity between multiple regions of interest [18, 19], thus providing a lens into the heterogeneity of neural structure within and across different brain areas.

Here, we introduce a three-dimensional neuroanatomical dataset extracted from a validated, in-vitro mouse thalamo-cortical sample spanning six anatomically distinct regions of interest (somatosensory cortex, two thalamic nuclei, zona incerta, striatum and hypothalamus) [18]. This dataset was reconstructed using X-ray microtomography to reveal a diverse composition of microstructures (e.g., myelinated axons, cell bodies, and vasculature) within each region at isotropic, micron-scale resolution. For a selected number of regions, we also provide validated annotations performed at the area level (which identifies regions of interest) and at the pixel level (which identifies microstructures). To technically validate the dataset, human annotators assessed two series of extracted images. We found that both annotators were able to classify images from these datasets reliably and accurately, likely in part due to the heterogeneity of the microstructures throughout each regions of interest. In the spirit of open-access to the scientific community, this dataset and multi-scale annotations are available for download from an online, interactive atlas (http://bossdb.org/project/prasad2020). This atlas can be examined by any individual, without the requirement of an account or prior training.

Ultimately we envision this heterogeneous, 3D brain volume dataset as a resource not only for neuroscientists interested in exploring structures within this thalamocortical pathway, but also machine learning scientists seeking data diverse enough to test computer vision methods for brain area prediction and segmentation. We further believe the provision of this dataset will prompt collaborative opportunities for both experimentalists and theorists interested in exploring neural circuitry at the micron-level and beyond.

## Methods

### Sample preparation

All animal experiments were approved by the Institutional Animal Care and Use Committee (IACUC) at the University of Chicago. The thalamocortical sample for this dataset was obtained from an 8 week old, C57BL/6J female mouse. The animal was deeply anesthetized using Euthasol (60mg/kg), then transcardially perfused. Vasculature was first flushed with 0.1M cacodylate buffer, followed by primary fixatives paraformaldehyde (2%) and glutaraldehyde (2.5%) in 0.1M cacodylate buffer. The brain was dissected from the skull and then post-fixed for 48 hours. Following multiple rinses in 0.1M cacodylate buffer, the brain was sliced on a vibratome at a thickness of 450 um until the thalamocortical slice was obtained [18]. At this point, a 1.7 by 6.5 mm strip of tissue which preserves the pathway from somatosensory cortex to the ventral posterior thalamic nucleus was dissected (see Figure 1A,B; [18]). Prior to imaging, the total estimated volume of this sample was 5 mm^3^. The sample was further post-fixed in paraformaldehyde (2%) and glutaraldehyde (2.5%) for 2 hours at room temperature, rinsed three times with 0.1M cacodylate buffer, and stored overnight in 0.1M cacodylate buffer at 4° Celsius. The tissue was then embedded with heavy metals as described by [20]. The sample was initially stained with (2%) buffered osmium tetroxide for 1.5 hours at room temperature followed by (2.5%) potassium ferrocyanide for 1.5 hours at room temperature. After rinsing with water, tissue was incubated in (1%) filtered thiocarbohydrazide at 40 degrees Celsius for 45 minutes. Following rinsing with water, the tissue was stained with another round of (2%) unbuffered osmium tetroxide for 1.5 hours at room temperature. After another thorough rinse with water, the sample was stained with (1%) aqueous uranyl acetate overnight at 4 degrees Celsius and at 50 degrees Celsius for 2 hours. Following the final water rinse, tissue was stained with lead aspartate for 2 hours at 50 degrees Celsius. This was followed by dehydration through a series of graded ethanols and propylene oxide, and a gradual infiltration of the tissue with epon resin. The infiltrated sample was incubated in 100% epon resin overnight, before being placed in 1.5 mm cylindrical tubing with fresh 100% resin. The preparation was then cured in an oven at 60° Celsius for 48-72 hours.

**Figure 1:**
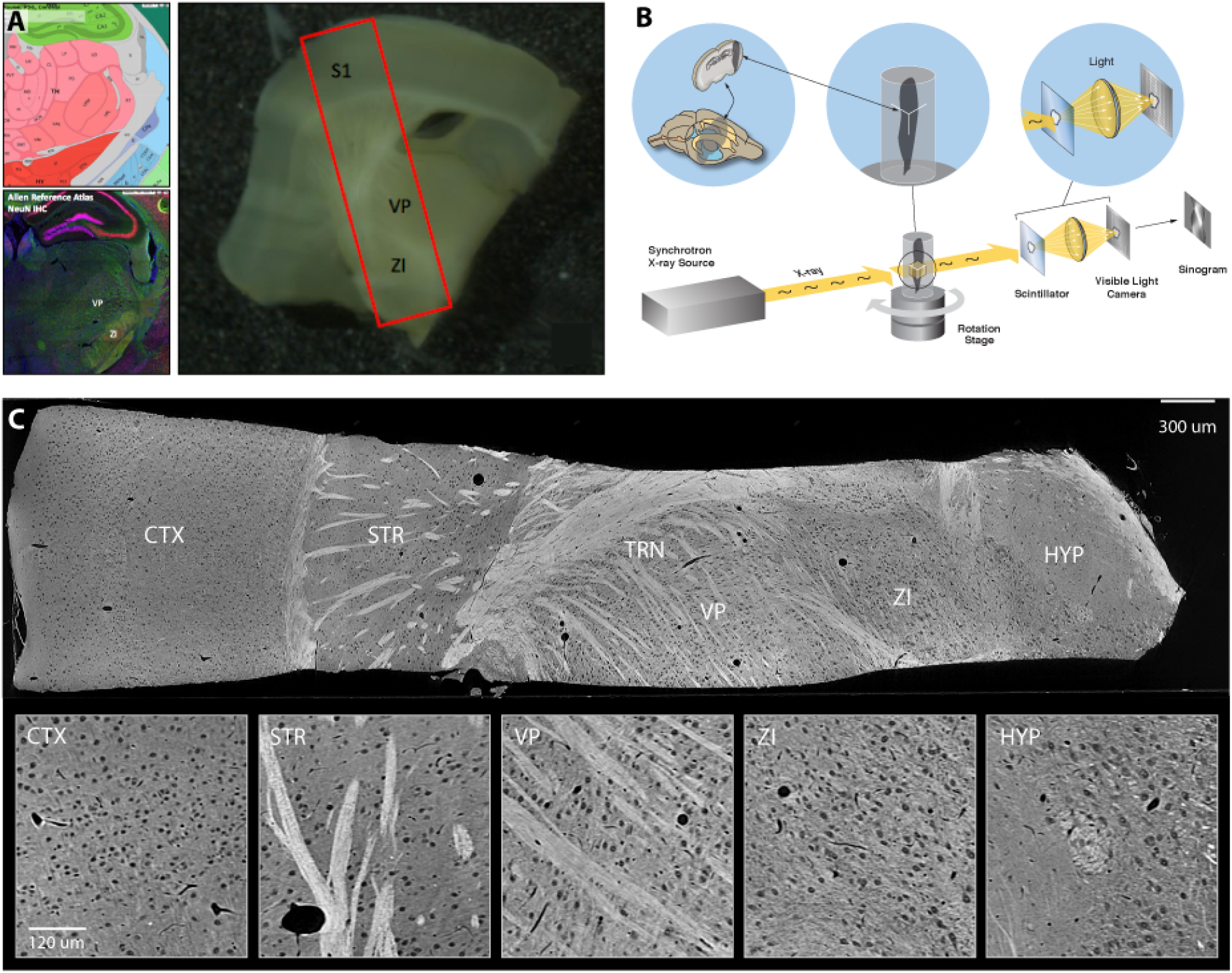
Overview of the thalamocortical sample cytoarchitecture and 3D reconstruction. Data from the Allen Reference Atlas (ARA) (http://mouse.brain-map.org/static/atlas) provides a schematic overview of the dataset regions of interest (A, top), along with cytoarchitectural differences as identified by NeuN staining (A, bottom) [10]. Photomicrograph to the right shows the thalamocortical slice used to obtain the dataset prior to dissection, outlined in red. S1 = somatosensory cortex; VP = ventral posterior nucleus; ZI = zona incerta. (B) Visualization of the synchrotron X-ray microtomographic data acquisition process. X-ray projections were acquired and reconstructed into a 3D image volume with 1.17 micron isotropic resolution. In (C), an example of a reconstructed image from the dataset following X-ray acquisition, highlighting the regions of interest visualized in the photomicrograph from A as well as others. Sub-panels below highlight the architectural diversity within and across regions of interest. From left to right: somatosensory cortex (CTX); striatum (STR); ventral posterior nucleus (VP); zona incerta (ZI); and hypothalamus (HYP).

### Imaging and reconstruction

Synchrotron X-ray tomography was performed on the embedded sample on the 32-ID beamline at the Advanced Photon Source in Argonne National Laboratory as described by Vescovi and colleagues [21]. X-ray radiographs were recorded with a detection system consisting of a LuAG:Ce scintillator converting X-rays into visible light that were magnified with a 5X objective lens onto a CMOS detector with 1920 × 1200 pixels (5.86 pixel size). To push the spatial resolution around 1 *μ*m, the detector was built with a large NA (0.21) long working distance Mitutoyo 5X objective lens with a resolving power of 1.3 *μ*m and 14 *μ*m depth of focus. To maintain its resolving power, the lens is coupled with a 13 *μ*m thick, thin film scintillator matching its depth of focus [22]. With a 5X magnification, the pixel size of each projection image was 1.17 *μ*m isotropic. Exposure for a single projection image took approximately 20ms, thus the total imaging acquisition time took approximately less than 60s per field-of-view.

Each single reconstructed dataset corresponded to a region of 1920 × 1920 × 1200 voxels. These volumes were then stitched together using Tomosaic software [21]. The entire volume was trimmed down to 720×1420×5805 voxels^3^ which corresponded to 0.842 × 1.661 × 6.792 mm^3^. Acquired data was stored in HDF5 files with Dxchange format (see [23]). To make subsequent analysis and labeling more aligned with an anatomical frame of reference, we virtually resliced the data to produce 720 images (each 1420×5805 pixels), where each image spans all cortical layers and traversed all sub-cortical regions of interest as well. This resulted in an image volume of 5.9 Gigavoxels total, including pixels outside of the sample. These images were chunked into 14 independent stacks comprised of 50.tiff files, spanning the entire length of the sample. These datasets were converted to 8 bits from 32 bit precision after histogram normalization.

### Ground truthing and annotation for brain areas (regions of interest)

6 regions of interest were annotated by an experienced neuroanatomist using a macroscale view of the entire thalamocortical sample (from somatosensory cortex to hypothalamus). 9 of the above 14 stacks were used for annotations, performed using the 3D segmentation software ITK-Snap [24]. Nine images (slices in Z) distributed uniformly throughout the volume (z = z1, z2,…) were annotated at the pixel level for each region of interest, approximately 50 images apart. The regions annotated included: somatosensory cortex (CTX), striatum (STR), the thalamic reticular nucleus (TRN), the ventral posterior (VP) nucleus of thalamus, zona incerta (ZI), the hypothalamus (HYP), and white matter (WM), which includes the internal capsule and corpus callosum. Each region’s annotation was therefore spaced 50 slices (58.5 *μ*m) apart throughout most of the volume (z = 109, 159, 209, 259, 309, 359, 409, 459), with the exception of the HYP which was only present for the first four stacks within the dataset. The span of the volume considered was bounded by slices 109 to 459, as regions beyond these slices were insufficiently visualized to provide complete annotations.

### Pixel-level annotations of selected regions of interest

We manually created ground truth pixel-level annotations of 4 major brain areas within the dataset: CTX, ZI, STR, and TRN. These regions were selected as they spanned the full extent of the sample, beyond simply cortical and thalamic regions of interest (i.e., through the inclusion of STR and ZI), making them well suited to studying architectural uniqueness within and across brain regions. To standardize the ground truth segmentations, we extracted 4 volumes of size (x:y:z) = (257:257:361), 1 for each of the 4 brain areas. The coordinates of each of these volumes are as follows, formatted as (xstart_xend ystart_yend zstart_zend): CTX (4600_4857 900_1157 110_471), ZI (1543_1800 650_907 110_471), STR (3700_3957 500_757 110_470), TRN (3063_3320 850_1107 110_471). Within each of these (257:257:361) brain area volumes, we densely annotated (starting at index z = 0) slice z = 30, 60, 90, 120, 150, 180, 210, 240, 270, 300, 330. This results in 11 densely annotated images per region and 44 images in total across the four areas.

### Data Records

We stored the 5.9 gigavoxel volume dataset and annotations in figshare [25] and in bossDB (http://bossdb.org/project/prasad2020). bossDB is a spatial database that is optimized for access and visualization of three-dimensional neuroimaging data, allowing users to efficiently and dynamically access different sub-volumes of the data [26]. bossDB connects seamlessly to Neuroglancer [27] which enables interactive visualization of large-scale, volumetric images, annotations, and analytics results from a web browser. Within this web portal, users can view and navigate the data in three dimensions using public access credentials (no account creation required). In addition to these web-based tools, we provide example Jupyter notebooks to demonstrate approaches for downloading raw data and annotations, and applying simple analysis algorithms to subvolumes of data (http://nerdslab.github.io/xray-thc). These tools directly address the needs of both novice and expert users and are easily adapted to additional use-cases.

The stored data and annotations consist of the following three sets of images and annotations:

- *Original Image Data* [25]: The raw images are rescaled from 32-bits and stored in 8-bit format, and stacked into an image volume of size 5805×1420×720 (x,y,z). The resolution is 1.17 *μ*m isotropic. Valid labels span the entire X, Y, and Z axis of the volume. Slices spanning from z=15 to z=670 contain brain specimen, and outside of this range, the epon block the sample is embedded in is visible. In bossDB, these annotations are stored in a channel called images.
- *Region-of-interest (ROI) Annotations* [28]: manually labeled pixel-level annotation of brain regions of interest. Valid labels span the entire X and Y axis of the volume. Valid z slices are 109, 159, 209, 259, 309, 359, 409, 459. The labels are 0-> no label; 1-> cortex; 2-> striatum; 3-> trn; 4-> vp; 5-> zona incerta; 6-> internal capsule; 7-> hypothalamus; 8-> corpus callosum. In bossDB, these annotations are stored in a channel called `region_of_interest`.
- *Pixel-level Microstructure Annotations* [29]: manually labeled pixel-level annotation of neural microstructure. Four volumes of size 361×257×257 are annotated. The first volume, from cortex, spans z from 110 to 471, y from 900 to 1157, and x from 4600 to 4857. The second volume, from Striatum, spans z from 110 to 471, y from 500 to 757, and x from 3700 to 3957. The third volume, from VP, spans z from 110 to 471, y from 850 to 1107, and x from 3063 to 3320. The fourth volume, from Zona Incerta, spans z from 110 to 471, y from 650 to 907, and x from 1543 to 1800. The labels are 0-> no label (background); 1-> vasculature; 2-> cell body; 3-> myelinated axon. In bossDB, these annotations are stored in a channel called `pixel_annotation`.

### Technical Validation

#### Examination of pixel-level features across regions of interest

As one of the defining features of this thalamocortical dataset is it’s diverse collection of brain regions, we sought to assess the quality of cytoarchitectonic heterogeneity by examining 4 major brain areas at a microstructural level (CTX, ZI, STR, TRN). Specifically, each structure’s anatomical composition (cells, blood vessels, and axons) as well as the intensities of pixels within each region (see Figure 2A,B) were measured. We determined the composition of features for each region (assessing the percentage of blood vessels, cells and myelinated axons within each), based on manual annotations over 128 images (32/class, 150 *μ*m by 150 *μ*m in size). While relatively similar numbers of cells and vasculature were identified within each region of interest, the fraction of axons annotated within each region varied dramatically, with far fewer axons identified in the CTX and ZI relative to a high proportion of axons in the STR and TRN (Figure 2C). Corroborating this finding are manual annotations which confirmed TRN and CTX were highly dissimilar in their microstructural composition, whereas the TRN and STR were highly similar as measured by the Kullback–Leibler (KL) divergence (Figure 2E). The KL divergence between pixel intensity distributions (Figure 2D) revealed that CTX is most dissimilar from TRN, which is evident given the vast differences in microstructure between these two regions of interest as described above. One unexpected finding was, in spite of the difference in their microstructural composition, STR and CTX were highly similar in their pixel intensity distributions. The 3D reconstruction of our thalamocortical dataset with microCT thus provided richer details that are unobservable upon examining stained, thinly sliced tissue using traditional light microscopy methods.

**Figure 2:**
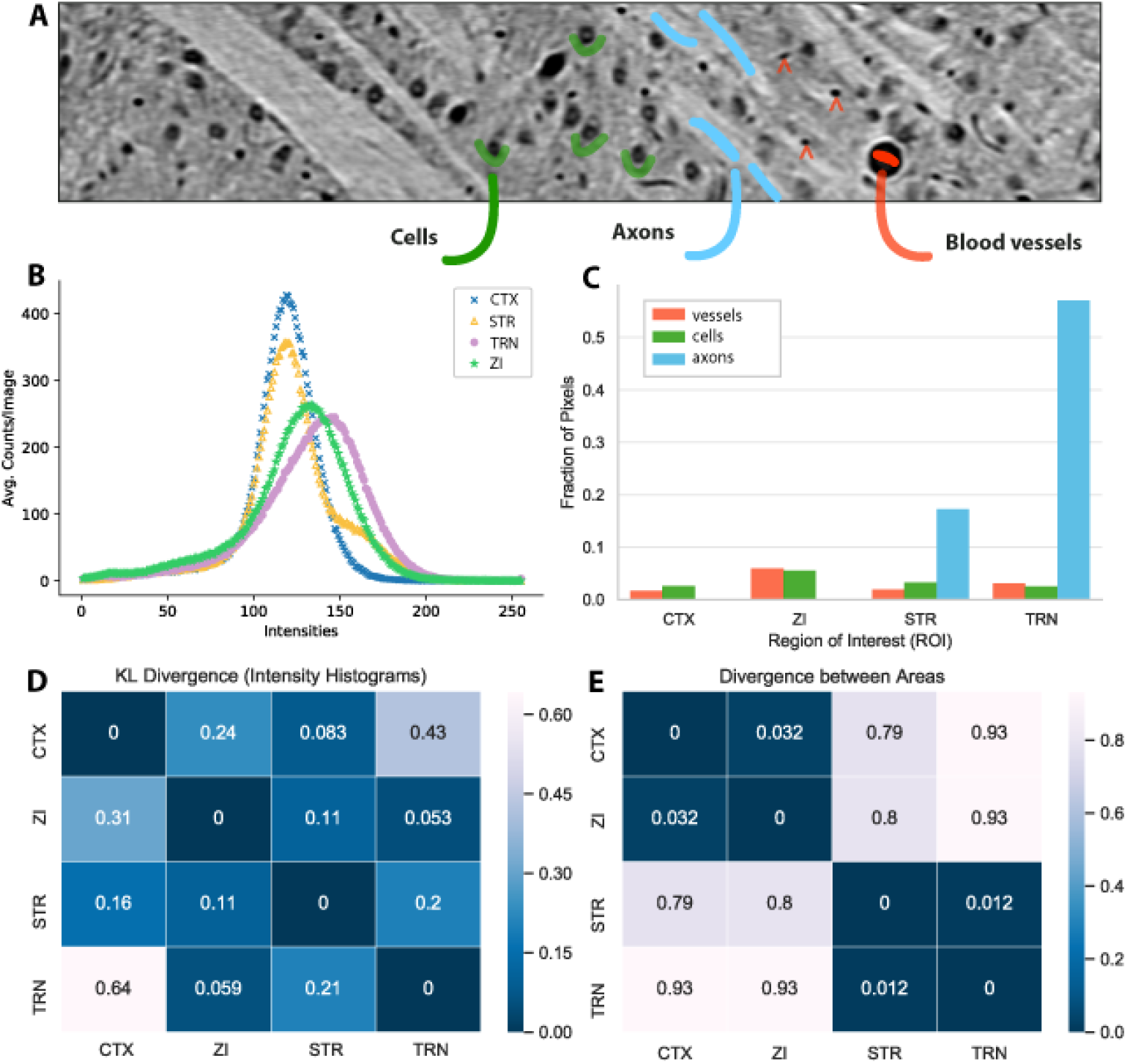
Validation of neuroanatomical heterogeneity within the dataset. (A) Example of microstructures identified within the thalamus, including cells, axons and blood vessels. (B) The distribution of pixel intensities across four selected regions of interest within the dataset (CTX, STR, TRN, ZI). (C) The distribution of pixels classified by underlying microstructure class (cell, blood vessel, axon) within each region of interest. In (D-E), we show the KL-divergence between: the intensity distributions across the selected regions (D), and the microstructural composition of selected regions as measured with dense manual annotations (E).

#### Region of interest prediction from local views of the microarchitecture

This dataset is comprised of a range of brain regions which can be visualized and annotated by any user interested in exploring macro-level or local characteristics. To validate the use of this dataset with annotators, we assessed whether humans can accurately predict regions of interest within the sample using only a small field-of-view (150×150 microns); see Figure 3A. For simplicity, TRN and VP were combined into a single region of interest (VP). The internal capsule and corpus callosum fiber tracts were included in this study, and categorized as WM. We then provided a training set of 48 images (8 per region of interest) for 2 annotators to examine. After studying these examples, each annotator was provided with a test set of 180 novel images (30 per region of interest) to sort into one of six region of interest categories.These annotators had previously studied example images from each region of interest prior to classifying these images, but were not as extensively trained as that of an experienced neuroanatomist. Interestingly, both annotators classified images within the CTX, STR and WM to a high degree of accuracy(>80%), whereas images from HYP, VP and ZI proved more challenging to classify (see Figure 3B). Generally, CTX images were classified to the same degree of accuracy as WM images relative to other regions of interest (see Figure 3C). We also noted that the annotators themselves varied in their classification performance; while Annotator 2 was slightly more accurate at classifying images from CTX relative to Annotator 1, they both had significantly more difficulty with identifying images from the HYP (*p < 0.05). It was also evident that the images from HYP were most challenging to classify, regardless of annotator and relative to those from CTX and STR (which had highly distinguishing microstructures, and therefore more likely to be correctly classified). Collectively, these findings support this heterogeneous 3D imaging dataset serving as a generalizable and useful resource for the field, given that human annotators can use it to a high degree of success for image classification.

**Figure 3:**
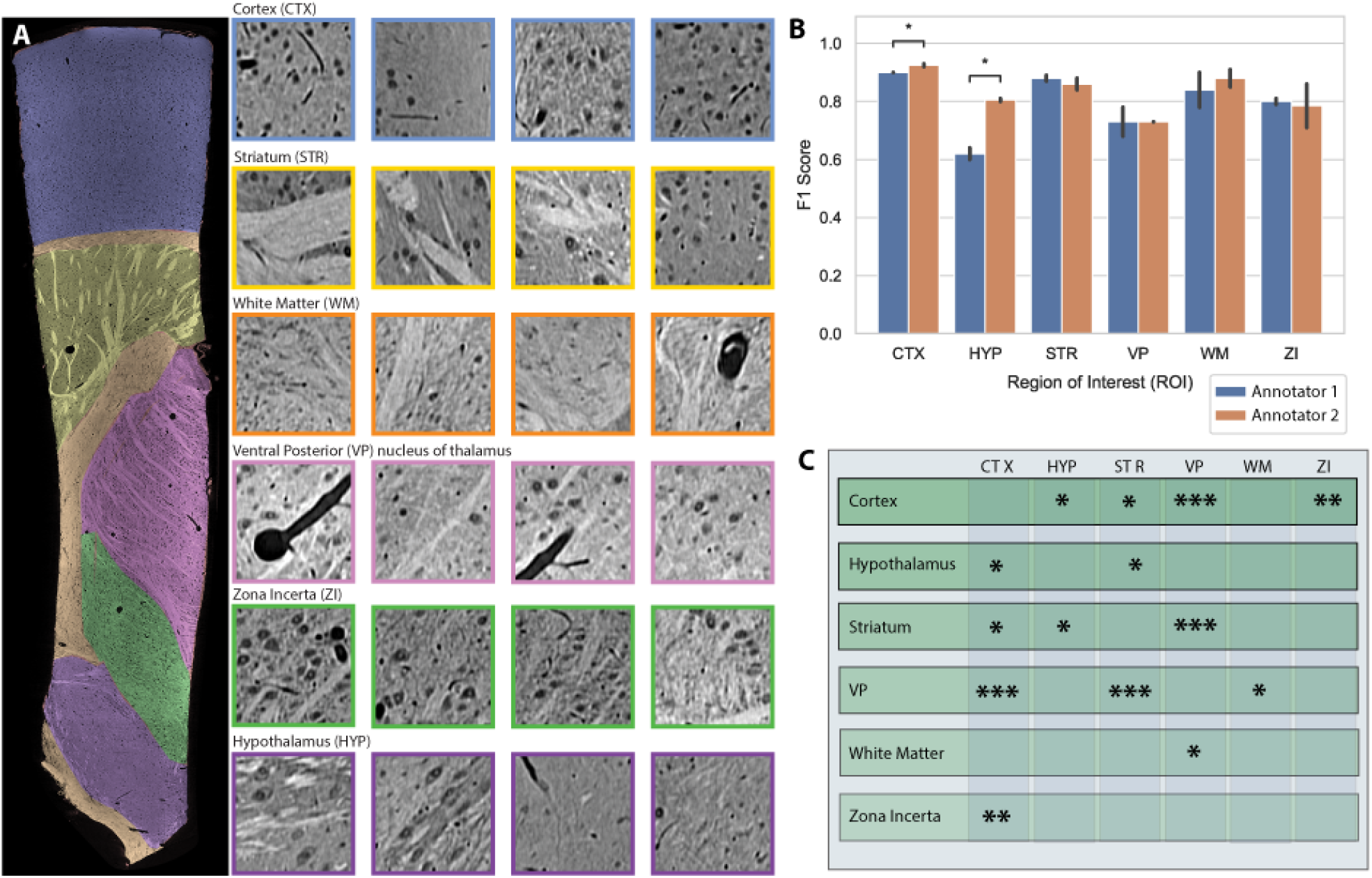
Brain area prediction performance. (A) An annotated image from the dataset with each region of interest overlaid as a distinct color (left) and 150×150 micron snapshots from within (right), highlighting microstructural heterogeneity within each region. (B) The performance (f1-score) and inter-rater reliability of two annotators classifying image patches similar to those visualized in A. Both annotators classified images into one of six different brain areas; each test set consisted of 180 images (30/class) for a total of 360 images classified. (C) Summary of significance in annotators’ ability to accurately predict a region of interest relative to others. Asterisks denote prediction measures that are significantly different, where * is used to denote p < 0.05, ** denotes p < 0.01, and *** denotes p < .001).

## Code Availability

Code for downloading the data and annotations in bossDB can be found in the ‘data_access_notebooks’ folder here: https://github.com/nerdslab/xray-thc. A Jupyter notebook for generating the results in Figures 2–3 can be found in the ‘analysis_notebooks’ folder in the same repo; this analysis notebook, datasets, and images used for the inter-rater reliability study, are also provided through figshare to facilitate reproducibility [30]. These examples are written in Python 3 and executed using Jupyter notebooks, a cross platform Python solution.

## Acknowledgements

E.L.D, A.H.B, and J.D.M were supported by award NSF IIS-1755871. E.L.D., E.J, and W.G.R were supported by NIMH grant R24MH114799. Support for bossDB comes from NIMH grant R24MH114785. M.D and C.J were supported by NIMH awards U01 MH109100 and R01 MH115265. This research used resources of the Advanced Photon Source, a U.S. Department of Energy (DOE) Office of Science User Facility operated for the DOE Office of Science by Argonne National Laboratory under Contract No. DE-AC02-06CH11357. We thank Michelle Alba for her assistance with the brain area prediction experiments described above.

## Author Contributions

J.A.P, E.L.D, K.P.K and N.K. conceived of the dataset. J.A.P and E.L.D designed the research and wrote the paper in collaboration with all co-authors. V.S. perfused the animal and processed the tissue for imaging; J.A.P dissected the thalamocortical sample.

V.D.A, K.F, M.D, R.V, C.J, D.G collected and/or generated the X-ray imaging data. V.D.A and K.F. created the custom imaging setup for microCT used to collect this data at beamline 32-ID at the Advanced Photon Source.

J.A.P performed annotation of all area-level ROIs. J.D.M provided all pixel-level annotations, curated the ground truth datasets and image volumes, and helped develop code to access subvolumes and annotations using the Bossdb API.

A.H.B conducted all aspects of the human annotation experiment (including the extraction of training datasets and analysis of annotator performance), analyzed and validated the data. E.C.J and W.G.R ingested data into bossDB and designed code to pull down and interact with data in bossDB and Neuroglancer.

